# Lipid-protecting disulfide bridges are the missing molecular link between ApoE4 and sporadic Alzheimer’s disease in humans

**DOI:** 10.1101/2025.01.17.633633

**Authors:** Christopher E. Ramsden, Roy G. Cutler, Xiufeng Li, Gregory S. Keyes

## Abstract

As the principal lipid transporter in the human brain, apolipoprotein E (ApoE) is tasked with the transport and protection of highly vulnerable lipids required to support and remodel neuronal membranes, in a process that is dependent on ApoE receptors. Human *APOE* allele variants that encode proteins differing only in the number of cysteine (Cys)-to-arginine (Arg) exchanges (ApoE2 [2 Cys], ApoE3 [1 Cys], ApoE4 [0 Cys]) comprise the strongest genetic risk factor for sporadic Alzheimer’s disease (AD); however, the *specific* molecular feature(s) and resultant mechanisms that underlie these isoform-dependent effects are unknown. One signature feature of Cys is the capacity to form disulfide (Cys-Cys) bridges, which are required to form disulfide bridge-linked dimers and multimers. Here we propose the overarching hypothesis that the super-ability (for ApoE2), intermediate ability (for ApoE3) or inability (for ApoE4) to form lipid-protecting intermolecular disulfide bridges, is the central molecular determinant accounting for the disparate effects of *APOE* alleles on AD risk and amyloid-β and Tau pathologies in humans. We posit that presence and abundance of Cys in human ApoE3 and ApoE2 respectively, conceal and protect vulnerable lipids transported by ApoE from peroxidation by enabling formation of ApoE homo-dimers/multimers and heteromeric ApoE complexes such as ApoE-ApoJ and ApoE-ApoD. We thus propose that the inability to form intermolecular disulfide bridges makes ApoE4-containing lipoproteins uniquely vulnerable to peroxidation and its downstream consequences. Consistent with our model, we found that brain-enriched polyunsaturated fatty acid-containing phospholipids induce disulfide-dependent dimerization and multimerization of ApoE3 and ApoE2 (but not ApoE4). By contrast, incubation with the peroxidation-resistant lipid DMPC or cholesterol alone had minimal effects on dimerization. These novel concepts and findings are integrated into our unifying model implicating peroxidation of ApoE-containing lipoproteins, with consequent ApoE receptor-ligand disruption, as the initiating molecular events that ultimately lead to AD in humans.

**Highlights:**

- *APOE* alleles are the strongest genetic risk factor for sporadic Alzheimer’s disease (AD)
- *APOE* alleles encode proteins that differ only in the number of Cys⟶Arg exchanges
- Despite 30 years of inquiry, mechanisms linking Cys⟶Arg exchanges to AD remain unknown
- PUFA-phospholipids induced disulfide bridge formation in ApoE3 and ApoE2 (but not ApoE4)
- We hypothesize that disulfide bridges in ApoE protect vulnerable lipids from peroxidation
- We propose that lipid-protecting disulfide bridges explain *APOE* allele-dependent AD risks

## 1. Introduction

As the principal lipid transporter in the human brain, apolipoprotein E (ApoE) is tasked with the transport and protection of essential—but highly vulnerable—lipids that are required to support and remodel the membranes of an estimated 86 billion neurons and 100 trillion synapses,[1-4] in a process that is dependent on neuronal ApoE receptors. The human genome includes three major *APOE* alleles that encode proteins differing only in the number of cysteine (Cys)⟶arginine (Arg) exchanges (ApoE2 [2 Cys, 33 Arg]), ApoE3 [1 Cys, 34 Arg], ApoE4 [0 Cys, 35 Arg]) (**Fig 1**). Remarkably, these seemingly innocuous exchanges have a profound impact on risk of developing sporadic, late-onset Alzheimer’s disease (AD),[5-9] (**Fig 1**) as well as the extent of AD-defining amyloid-β (Aβ) and hyper-phosphorylated Tau (pTau) containing lesions.[9, 10]

**Fig 1.**
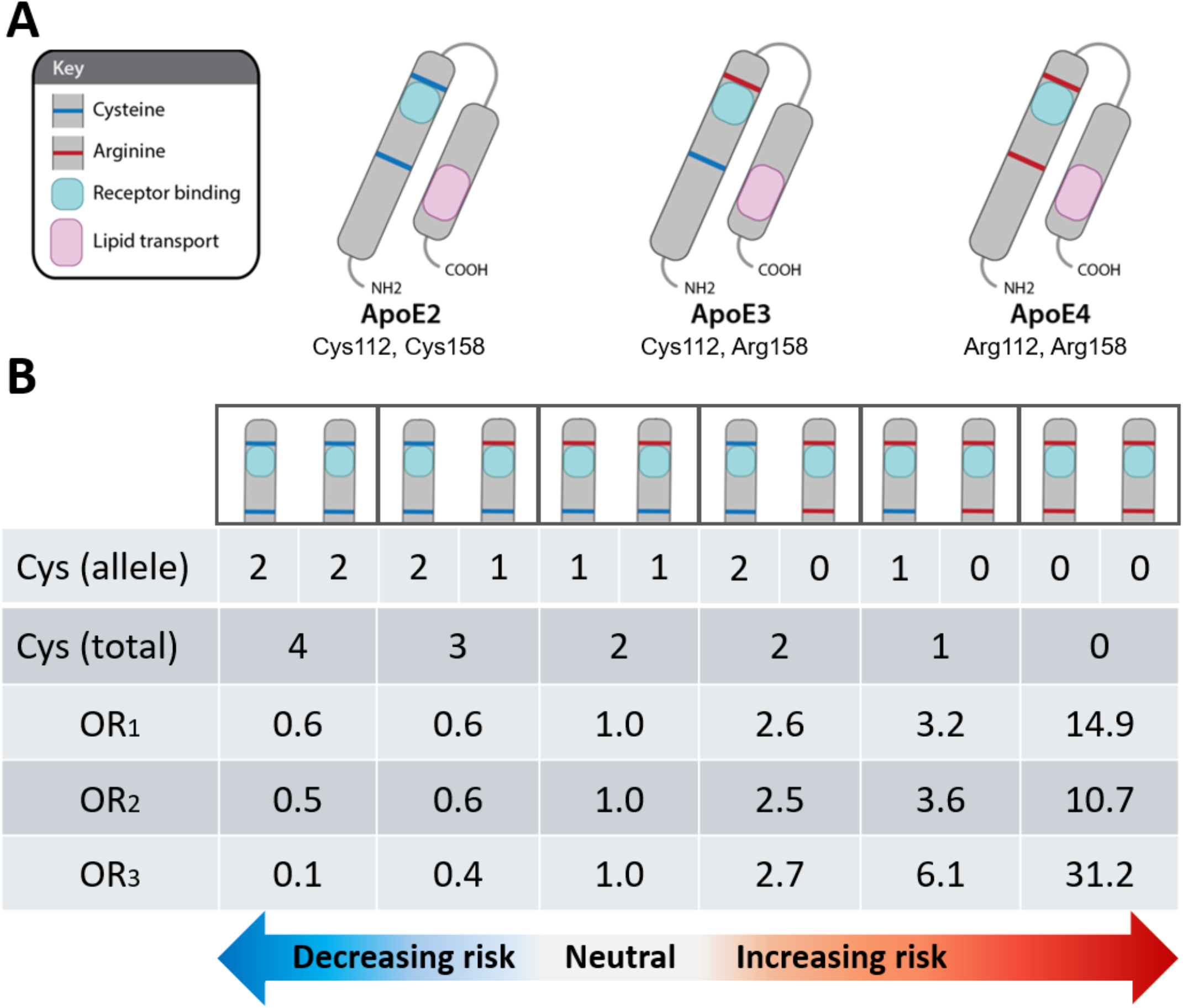
The APOE allele-dependent number of Cys residues predicts sporadic AD risk in humans. (**A**) The three major APOE alleles encode proteins differing only in the number of Cys⟶Arg exchanges (ApoE2 [2 Cys]), ApoE3 [1 Cys], ApoE4 [0 Cys]). (**B**) There is a strong, dose-dependent inverse relationship between the total number of Cys residues provided by two alleles (allelic dose) and AD risk, with the lowest risk in APOE2 homozygotes (4 Cys), intermediate risk in APOE3 homozygotes (2 Cys, referent), and the highest risk in APOE4 homozygotes (0 Cys). OR_1_ indicates the odds ratios reported by Farrer et al. (1997); OR_2_ and OR_3_ indicate the odds ratios reported by Reiman et al. (2020) for neuropathologically unconfirmed and neuropathologically confirmed AD cases, respectively. APOE3 homozygosity, the most common genotype, was considered the neutral referent for genotypic comparisons.

The *APOE2* allele (with two Cys at positions 112 and 158 of the mature protein [**Fig 1**]) is the strongest genetic protective risk factor for AD,[7, 11, 12] and is associated with the fewest Aβ plaques and pTau tangles.[10] The *APOE3* allele (with one Cys at position 112 [**Fig 1**]) has moderate AD risk and intermediate levels of Aβ plaques and pTau tangles.[9] The *APOE4* allele (with zero Cys [**Fig 1**]) is the strongest genetic risk factor for AD,[9] and has earlier onset,[6, 11] a more aggressive clinical course, and more extensive and severe Aβ and pTau pathologies.[10] ApoE accumulates together with its sister apolipoprotein (ApoJ, also known as clusterin) in the central core of Aβ plaques surrounded by pTau-enriched neurites known as neuritic plaques,[13-16] placing ApoE and ApoJ at the interface of hallmark Aβ and pTau lesions. ApoE and ApoJ also accumulate together with Aβ in cerebral amyloid angiopathy lesions.[17] The human brain and glia-derived brain lipoproteins are enriched in dozens of polyunsaturated fatty acid (PUFA)-containing lipid species [2-4] that are highly vulnerable to peroxidation;[18] extensive brain lipid peroxidation has been reported even in the earliest stages of mild cognitive impairment (MCI) and AD [19, 20] (reviewed in [15]).

Despite more than three decades of evidence supporting the dominant role of *APOE4* and *APOE2* genetics, the enrichment of ApoE protein in hallmark lesions, and extensive lipid peroxidation in AD brains, *specific* molecular feature(s) and unambiguous feature-dependent mechanism(s) directly linking Cys⟶Arg exchanges in ApoE to AD risk have not been clearly defined. ApoE is a multi-functional protein;[21] this multiplicity of function may have paradoxically hindered identification of the most salient molecular feature(s) and mechanism(s) that drive AD risk. For example, a team of AD experts reviewed extensive published evidence linking ApoE to AD and proposed an ‘ApoE Cascade Hypothesis’[22] wherein numerous ‘biochemical and biophysical properties, including ApoE structure, lipidation, protein levels, receptor binding, and oligomerization’[22] impact a disease cascade that ultimately leads to AD. Importantly, however, they did not endorse one upstream or overarching molecular feature or feature-dependent mechanism of ApoE as the principal driver of this risk.[22] Other investigators proposed that molecular features specific to ApoE4, such as unique salt bridge interactions [23, 24] or increased tendency to aggregate,[25-28] could account for increased risks in *APOE4* carriers. However, because ApoE4-specific features do not readily explain the protective effects of *APOE2* (**Fig 1**), these hypotheses do not provide one central mechanism to explain the full spectrum of *APOE* variants on AD risk.

The aim of this hypothesis paper is to propose one specific, overarching and unambiguous molecular feature and feature-dependent mechanism that: (1) can explain the full spectrum of common *APOE* variants on AD risk (including protective effects of *APOE2* and harmful effects of *APOE4*); (2) is traceable directly back to the Cys⟶Arg exchange(s) that are the fundamental difference between *APOE* alleles; (3) can explain both loss of function and gain of toxic function aspects of ApoE4 biology; and (4) can be readily integrated into a coherent model with other hallmark and emerging genetic and neuropathological features of AD.

## 2. The hypothesis

One signature feature of Cys is the capacity to form disulfide (Cys-Cys) bridges,[29] which in turn are required to form disulfide-linked protein dimers and multimers (**Fig 2**). Here, we propose the overarching hypothesis that the super-ability (for ApoE2), intermediate ability (for ApoE3) or complete inability (for ApoE4) to form disulfide bridge-linked ApoE multimers or dimers is the central molecular determinant accounting for the disparate effects of *APOE2, APOE3* and *APOE4* alleles on Aβ and pTau pathology and AD risk in humans. We further propose that the intermolecular disulfide bridges present in ApoE2 multimers (**Fig 2A**) and ApoE3 dimers (**Fig 2B**) decrease AD risk specifically by concealing and protecting vulnerable lipid cargo transported by ApoE-containing lipoproteins from peroxidation, thus preventing the deleterious downstream consequences of lipoprotein peroxidation (reviewed in [15]). We therefore propose that the absence of Cys, and resulting inability to form intermolecular disulfide bridges (**Fig 2C**), make ApoE4-containing lipoproteins uniquely vulnerable to peroxidation (**Fig 2D-F**) and ensuing peroxidation-induced pathological events, including the aldehydic adduction of Lys and His residues within the receptor binding motif of ApoE and crosslinking of ApoE with ApoE receptors, initiating a disease cascade that ultimately manifests as AD (**Fig 2G-H**).

**Fig 2.**
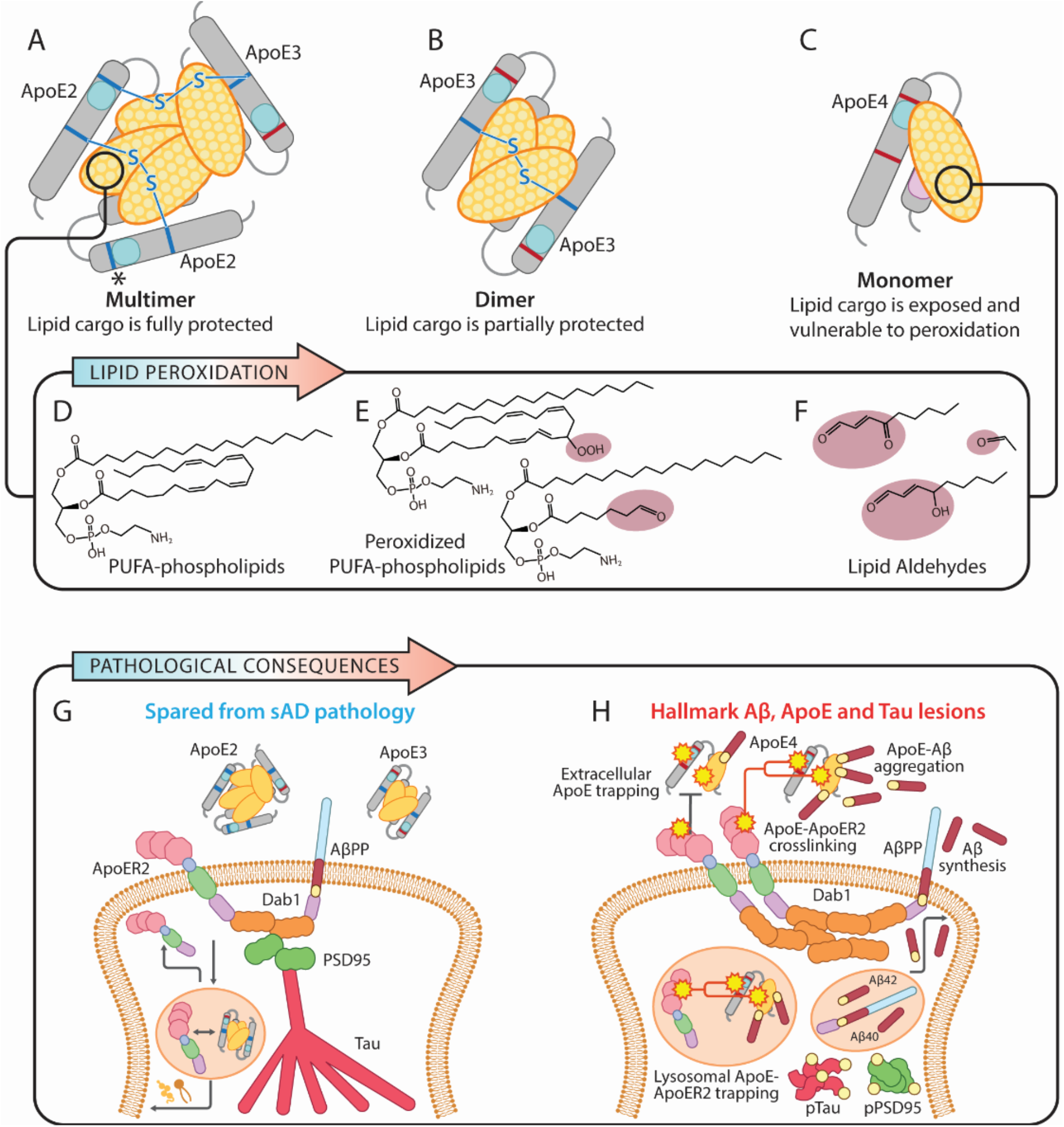
ApoE disulfide bridge-dependent lipoprotein protection model of AD. According to our hypothesis: (1) the peroxidation of vulnerable lipids transported by ApoE initiates a disease cascade that ultimately manifests as AD; and (2) disulfide bridge-dependent ApoE dimer/multimerization decreases AD risk by protecting its lipid cargo from peroxidation and its deleterious molecular consequences. Cys residues present in ApoE2 and ApoE3 are required to generate disulfide bridges (depicted as blue lines in **A** and **B**, not to scale), which connect ApoE monomers into dimers and multimers. ApoE contains a hydrophobic domain that transports cholesterol and highly vulnerable polyunsaturated (PUFA)-containing phospholipids. In this model, disulfide-dependent multimerization of ApoE2 generates a hydrophobic core between ApoE proteins that conceals and protects lipid cargo from peroxidation (**A**). Disulfide-dependent dimerization of ApoE3 partially conceals and protects lipid cargo from peroxidation (**B**). Absence of disulfide bridge-dependent dimerization in ApoE4 (**C**) leaves brain-enriched PUFA-phospholipids (**D**) exposed and vulnerable to peroxidation (**E**). Resulting lipid aldehydes (**F**) trigger pathological protein adduction and crosslinking. Panel **G** depicts physiological conditions that predominate in the absence of APOE4, including functional ApoE-ApoE receptor interactions and internalization, ApoE receptor recycling, limited AβPP cleavage, and suppressed Tau phosphorylation. Panel **H** depicts pathological conditions in APOE4 carriers wherein peroxidation of ApoE4-containing lipoproteins disrupts ApoE-ApoER2 interactions, traps peroxidized ApoE-containing lipoparticles in the extracellular space where they facilitate Aβ polymerization, disrupts lysosomal function, and increases Aβ synthesis and Tau phosphorylation. These pathological events are proposed to underlie hallmark Aβ plaques, Tau tangles and emerging pathological features of AD. The asterisk (*) in **A** indicates that free Cys residues in ApoE2-containing multimers can form intermolecular disulfide bridges with additional ApoE monomers (not shown), or with other Cys-containing proteins to form heteromeric complexes (see **Fig 3**).

In its simplest form, this hypothesis focuses on how disulfide bridge-linked ApoE3 and ApoE2 homo-dimers and multimers conceal and protect vulnerable lipids (**Fig 2**); however, the presence of Cys also endows ApoE2 and ApoE3 with the ability to form disulfide-linked heteromeric complexes via unpaired Cys residues present in other human proteins (indicated by * in **Fig 2A**). For example, disulfide-linked ApoE3-ApoAII complexes are known to be abundant in serum lipoproteins.[30-34] Yamauchi et al [35] showed that serum-derived ApoE3-ApoAII complexes can inhibit irreversible Cys amino acid oxidation induced by H_2_O_2_, suggesting that ApoAII could exert disease-modifying effects in peripheral tissues. Importantly, however, ApoAII is selectively expressed by liver [36] and immunohistochemical studies indicate that ApoAII is not detectable in human brain (except for brain vasculature).[37] Moreover, ApoAII does not appear to be expressed by astrocytes,[36] which secrete the majority of ApoE particles in the human brain.[38-40] Astrocytes and other glia do secrete large amounts of ApoJ [39, 40] and ApoD,[41] which have established roles in lipid transport and detoxification.[42, 43] We therefore consider these two apolipoproteins to be prime candidates for inclusion into disulfide-linked heteromeric complexes within ApoE-containing brain lipoproteins.

The canonical ApoJ proteoform encoded by CLU isoform 1 (referred to here as ApoJ_1_) contains ten Cys residues[36]; however, since all ten are entwined in intermolecular disulfide bridges connecting alpha and beta chains, there are no free Cys residues available for disulfide linkage with ApoE2 or ApoE3 (**Fig 3A**). Intriguingly, however, a comparatively understudied isoform (CLU Isoform 2), which is reported to be selectively overexpressed in individuals carrying the AD-protective minor rs11136000T allele,[44] encodes an ApoJ proteoform containing three additional Cys residues in the extreme N-terminus [36] (referred to here as ApoJ_2_) (**Fig 3B**). We posit that these free Cys residues endow ApoJ_2_ with the ability to conceal and protect vulnerable lipids by forming disulfide-linked heteromeric complexes with ApoE2 and ApoE3 (but not ApoE4), suggesting a novel and plausible explanation for the strong protective association between the rs11136000T allele and AD.[8] The finding by Moreno et al [45] that the strong protection afforded by the rs11136000T allele is limited to APOE4 non-carriers with no effect in APOE4 carriers, is consistent with our hypothesis wherein the protective effects of ApoJ_2_ are dependent on formation of disulfide-linked heteromeric ApoE-ApoJ_2_ complexes.

**Fig 3.**
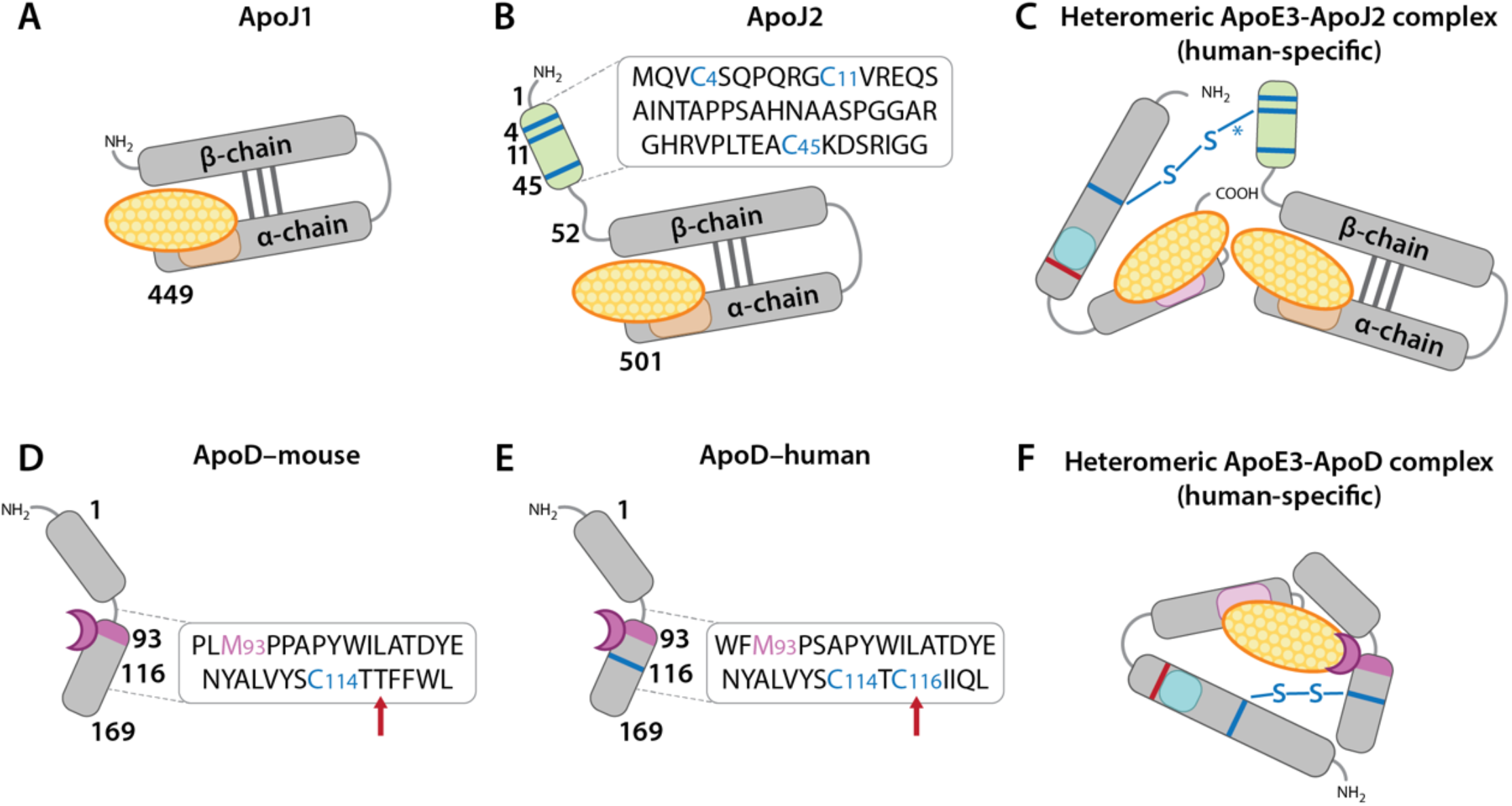
Putative disulfide bridge-linked heteromeric apolipoprotein complexes in human brain. According to this sub-hypothesis: (1) the formation of heteromeric disulfide-linked apolipoprotein complexes decreases AD risk by protecting vulnerable lipid cargo transported by ApoE-containing lipoproteins from peroxidation and its deleterious molecular consequences; and (2) free Cys residues endow ApoE2 and ApoE3 (but not ApoE4) with the ability to generate disulfide bridge-linked heteromeric complexes with other glia-derived apolipoproteins such as ApoJ and ApoD. The canonical CLU isoform 1 encodes an ApoJ protein lacking any free Cys (referred to here as ApoJ_1_) (**A**). By contrast, the understudied CLU isoform 2 encodes an ApoJ protein containing three additional Cys residues (referred to here as ApoJ_2_) (**B**). We posit that these extra Cys residues endow ApoJ_2_ with the unique ability to protect lipid cargo by forming disulfide bridge-linked heteromeric complexes with ApoE2 (not shown) and ApoE3 (**C**) but not ApoE4. The Cys residues in rodent ApoD are all entwined in intramolecular disulfide bridges (**D**). Human ApoD contains one additional unpaired Cys (116) that is not present in rodent models (see red arrows in **D-E**). We posit that this human-specific Cys116 endows human ApoD with the unique ability to form disulfide bridge-linked heteromeric complexes with ApoE2 (not shown) and ApoE3 (**F**) but not ApoE4. Further, we propose that lipid-protecting and peroxide reducing capabilities of ApoJ and ApoD endow these human-specific heteromeric complexes with the ability to suppress and neutralize the peroxidation of brain lipoproteins. The asterisk (*) in **3C** indicates that Cys112 in ApoE3 could form a disulfide bridge with Cys4, Cys11, or Cys45 in ApoJ_2_.

Like ApoJ, ApoD is a glia-secreted apolipoprotein that is present in ApoE-containing lipoproteins, and has increased expression in AD and in response to oxidative stress. [42, 43] The methionine 93 (Met93) endows ApoD with the ability to neutralize lipid peroxide radicals [42, 43] (**Fig 3**). Most of our current understanding of ApoD is extrapolated from rodent models wherein ApoD lacks unpaired Cys residues [36] that are needed to form disulfide-linked heteromers (**Fig 3C**). Intriguingly, however, human ApoD contains one unpaired Cys (Cys116) that is not present in rodents [36, 46] (see red arrows in **Fig 3D-E**). This free Cys may account for detection of disulfide-linked ApoD-ApoB complexes in human plasma lipoproteins reported back in 1989.[47] Here we posit that this human-specific Cys116 endows ApoD in human brain with the ability to form homologous disulfide-linked heteromeric complexes with lipoprotein-associated ApoE2, ApoE3, and ApoJ_2_ (but not ApoE4 or ApoJ_1_) (**Fig 3F**). Further, we propose that these putative human-specific ApoE2-ApoD and ApoJ_2_-ApoD complexes protect brain lipoproteins from peroxidation and its downstream consequences.

### Can ‘disulfide deficiency’ in ApoE4 bridge the gap between loss of function and gain of toxic function?

There is a long-standing debate about whether the increased AD risk associated with *APOE4* is due to a ‘loss of function’ or a ‘gain of function’.[48] The concept of Cys and disulfide deficiency, which assigns a loss of protective, physiological function (inability to form disulfide bridges) to ApoE4 that in turn predisposes to a gain of toxic function (accelerated production of toxic peroxidized lipids), offers a straightforward mechanism that can reconcile both interpretations of ApoE4 biology (**Fig 4**). By assigning partial loss of function to ApoE3 (intermediate ability to form disulfide bridges) with consequent partial gain of toxic function (incomplete lipid protection), this model is proposed to explain the full spectrum of genetically-driven AD risk. This mechanistic paradigm and proposed sequence of molecular events (**Fig 4**) is attractive because it integrates ApoE and ApoJ genetics and biology with many previously disjointed observations such as extensive lipid peroxidation in AD brain, and environmental triggers for lipid peroxidation that are known or suspected AD risk factors (reviewed in [15]), into one unifying explanation for human sporadic AD.

**Fig 4.**
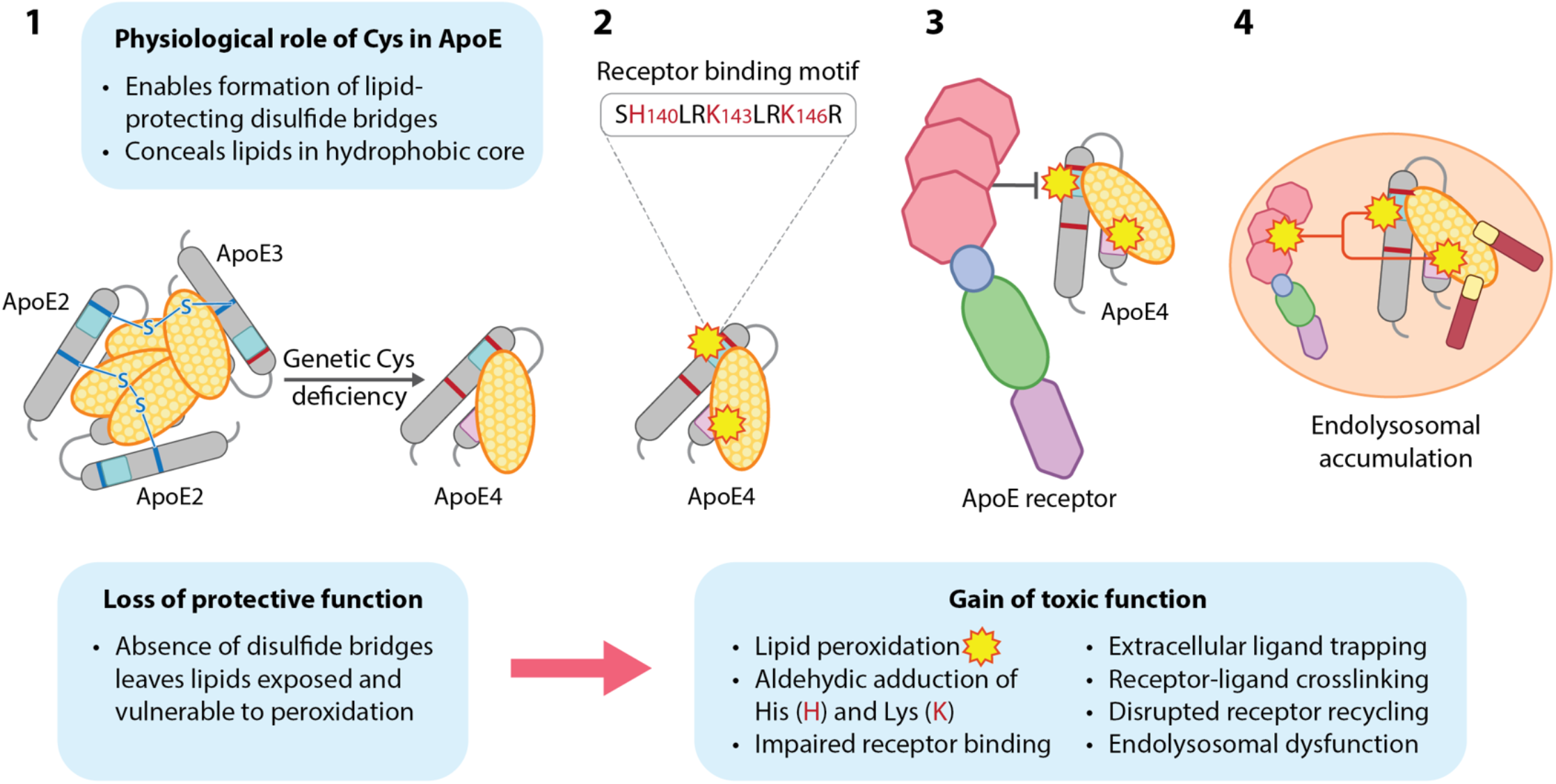
Disulfide deficiency in ApoE4: bridging the gap between loss of function and gain of toxic function. We propose a model wherein the genetically-determined lack of Cys in ApoE4 increases AD risk by precluding the formation of lipid-protecting intermolecular disulfide bridges. A 4-step model wherein this ‘loss of function’ leads to a ‘gain of (toxic) function’ by predisposing to lipid peroxidation is provided below. **Step 1**: Lack of disulfide bridges renders ApoE4 and its lipid cargo exposed and vulnerable to peroxidation and its downstream consequences. **Step 2**: Peroxidation of PUFA-containing lipids transported by ApoE generates reactive lipid aldehydes; consequent aldehydic adduction of K and H residues within the cationic receptor binding motif of ApoE disrupts physiological binding to ApoE receptors. **Step 3**: Disrupted ApoE ligand-receptor binding increases GSK3-mediated Tau phosphorylation and traps peroxidized ApoE-containing lipoproteins in the extracellular space, where they provide seeds for Aβ plaque formation (see **Fig 2H**). **Step 4**: Convergence of peroxidized ApoE-containing lipoproteins with reactive K and H residues in ApoE receptors such as ApoER2 leads to aldehyde-induced ApoE-ApoE receptor crosslinking. Internalization of peroxidized ApoE particles and crosslinked ApoE-ApoE receptor complexes compromises lipid delivery and ApoE receptor recycling. Crosslinked ApoE and ApoE receptors accumulate within enlarged, dysfunctional lysosomes.

We thus propose that the super-ability (for ApoE2), intermediate ability (for ApoE3) or complete inability (for ApoE4) to form lipid-protecting intermolecular disulfide bridges is the central molecular determinant accounting for the dose-dependent protective, neutral, and harmful effects of *APOE2, APOE3*, and *APOE4* alleles, respectively, on Aβ and pTau pathology and AD risk. This hypothesis is integrated into our unifying model [15, 16] implicating peroxidation of ApoE-containing lipoproteins, with consequent ApoE receptor-ligand disruption and compromised ApoE receptor 2-Disabled homolog-1 (ApoER2-Dab1) pathway function, as the initiating molecular events that ultimately lead to AD in humans.

## 3. Evaluation of the hypothesis

### 3.1. Do intact disulfide bridges protect PUFA-enriched ApoE lipoparticles from peroxidation?

PUFA-containing phospholipids—including phosphatidylethanolamine (PUFA-PE) species that are particularly vulnerable to peroxidation [18]—are key components of ApoE- and ApoJ-containing brain lipoproteins.[40] We posit that intermolecular disulfide bridges that are absent in ApoE4 monomers but present and abundant in ApoE3 and ApoE2 dimers and multimers, respectively (**Fig 2**), protect these vulnerable brain-enriched lipids from peroxidation. To generate reagents needed to answer this question, ApoE2, ApoE3, and ApoE4-containing lipoparticles can be synthesized by incubating recombinant ApoE with PUFA-containing, brain-enriched lipid species. Alternatively, ApoE-containing lipoproteins could be collected from the culture media of human *APOE2, APOE3*, and *APOE4*-expressing astrocytes. Crucially, to date the synthetic ApoE lipoparticles used in seminal studies characterizing ApoE structure were generated with peroxidation-resistant phospholipids such as 1,2-Dimyristoyl-sn-glycero-3-phosphatidylcholine (DMPC-14:0/14:0), an artificial lipid that is not present in nature,[49] and did not include any of the peroxidation-sensitive PUFA-containing phospholipid species that are highly enriched in the brain.[50-57] Remarkably, most seminal studies characterizing ApoE structure and biology also treated ApoE lipoparticles with thiol-reducing agents such as dithiothreitol (DTT), tris(2-carboxyethyl) phosphine (TCEP) or β-mercaptoethanol,[50-54, 57, 58] which cleave intermolecular disulfide bridges within ApoE2 and ApoE3-containing lipoparticles.[29] Lack of PUFA-containing phospholipids and/or use of thiol-reducing agents precluded evaluation of our model wherein intact intermolecular disulfide bridges serve a protective function by concealing vulnerable lipids in a hydrophobic core within ApoE-containing lipoparticles (**Fig 2-4**).

### 3.2. PUFA-phospholipids promote disulfide-dependent dimer/multimerization of ApoE3 and ApoE2

Consistent with our model, we found that incubation of recombinant ApoE3 and ApoE2 (but not ApoE4) proteins with the brain-enriched PUFA-PEs 1-1(Z)-Octadecenyl-2-Docosahexaenoyl-sn-glycero-3-PE (PE-P18:0/22:6n-3), 1-Stearoyl-2-Docosahexaenoyl-sn-glycero-3-PE (PE-18:0/22:6n-3), or 1-Stearoyl-2-Docosatetraenoyl-sn-glycero-3-PE (PE-18:0/22:4n-6), alone or as a mixture in combination with free cholesterol, increased disulfide-linked dimerization and multimerization (**Figs 5-6**). Among these, PE-P18:0/22:6n-3—a DHA-containing ethanolamine plasmalogen that is highly vulnerable to peroxidation and depleted in AD brain [59]—had the most pronounced effects on ApoE3 and ApoE2 multimerization. By contrast, incubation with the artificial, peroxidation-resistant lipid DMPC or cholesterol alone had minimal effect on disulfide bridge formation. Our observation that PUFA-phospholipids in general, and PE-P18:0/22:6n-3 in particular, promote disulfide-bridge formation supports our hypothesis wherein the presence of Cys is an intrinsic protective feature that allows ApoE2 and ApoE3 (but not ApoE4) to undergo disulfide-dependent conformational changes that conceal and protect highly vulnerable PUFA-containing lipids within a hydrophobic core; however, future studies are required to confirm the specific structural effects of intermolecular disulfide bridges within ApoE3 and ApoE2-containing lipoproteins. These disulfide-inducing effects of PUFA-phospholipids were reversed by DTT (**Fig 5-6**), highlighting crucial limitations of previous studies that used thiol-reducing agents and/or peroxidation-resistant lipids for ApoE lipidation. Future studies using additional brain-enriched PUFA-phospholipid species and more sophisticated models for ApoE lipidation with and without thiol-reducing agents, are needed to confirm these results and to fully understand effects of lipidation on disulfide bridge formation, lipoprotein peroxidation and downstream consequences, as described below.

**Fig 5.**
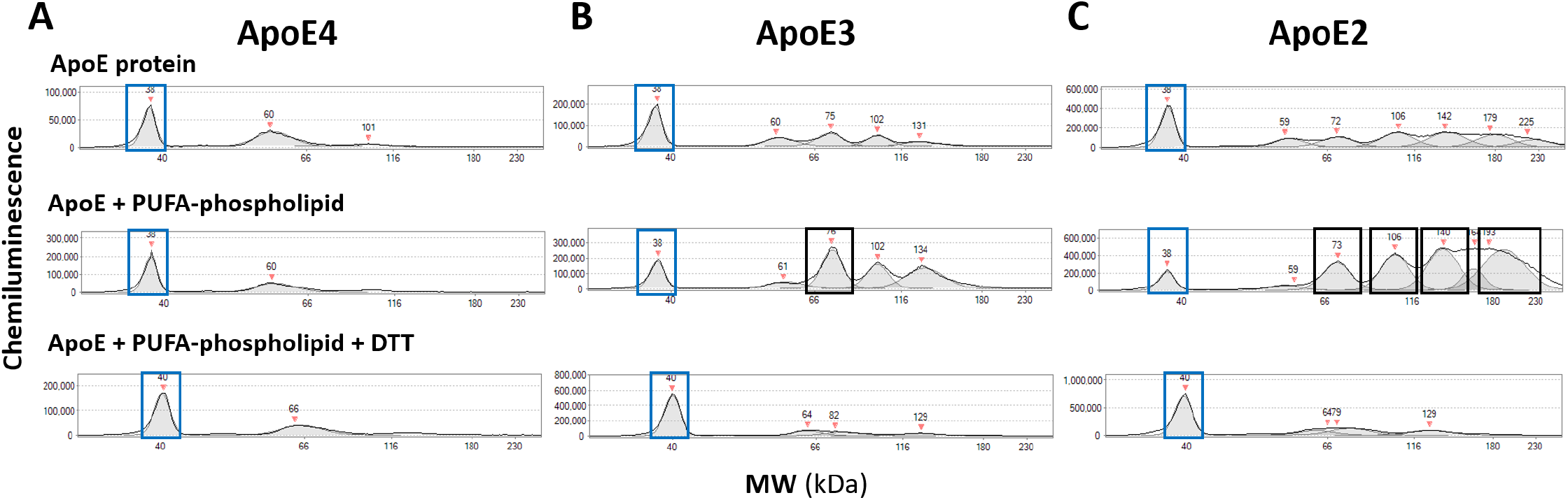
PUFA-phospholipids induce disulfide-dependent dimer/multimerization of ApoE3 and ApoE2. Recombinant ApoE4 (**A**), ApoE3 (**B**), and ApoE2 (**C**) proteins were incubated with the PUFA-phospholipid PE-18:0/22:4n-6 (see Suppl. methods). Protein separation and immunodetection were generated with the Jess™ automated capillary nano-immunoassay platform (Protein Simple, Bio-techne, 004-650) with chemiluminescence. Molecular mass marker (12-230 kDa), rabbit IgG anti-ApoE monoclonal (Abcam ab52607 [EP1374Y], 1 to 800 dilution) and anti-rabbit detection module (Novus DM-001) were loaded according to manufacturer instructions. Compass exposure setting 1 (1 second) was applied for visualization of peaks. Peaks corresponding to molecular weights of ApoE monomers are designated with blue rectangles; peaks corresponding to dimers and multimers are designated with black rectangles in B and C. Incubation with the PUFA-phospholipid PE-18:0/22:4n-6 had no apparent effect on ApoE4 dimerization (**A**). By contrast, incubation of ApoE3 (**B**) and ApoE2 (**C**) with PE-18:0/22:4n-6 produced prominent peaks with molecular weights corresponding to ApoE dimers and multimers, respectively. PE-18:0/22:4n-6 induced dimer/multimerization of ApoE3 and ApoE2 was reversed by addition of DTT. Capillary nano-immunoassay chromatograms are representative of experiments performed three times. Abbreviations: PUFA, polyunsaturated fatty acid; DTT, dithiothreitol.

**Fig 6.**
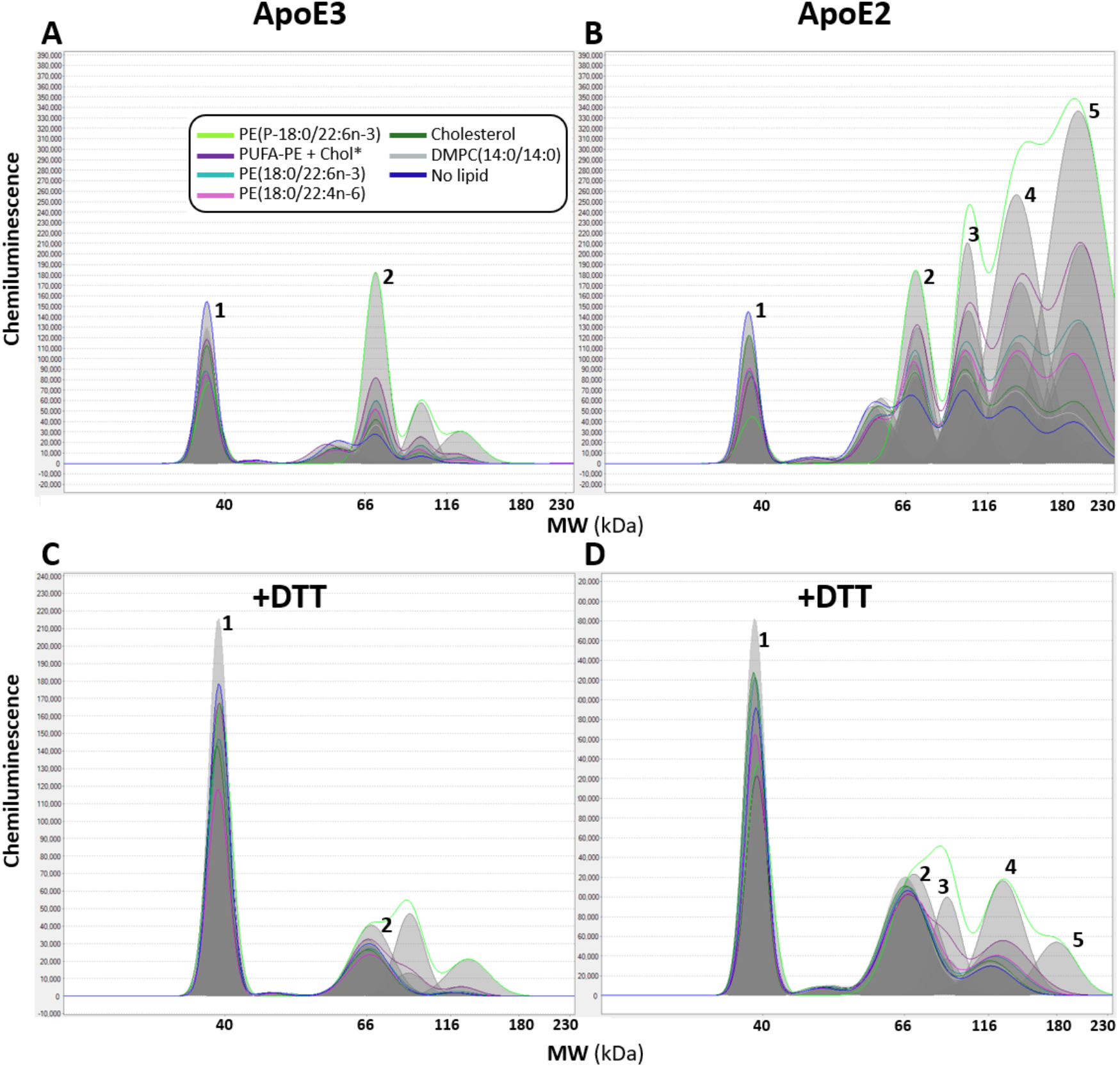
Cholesterol and DMPC have little or no effect on disulfide-dependent ApoE multimerization. Recombinant ApoE3 or ApoE2 proteins were incubated with equimolar concentrations of phospholipids, cholesterol, or a mixture thereof, without thiol-reducing agents (see Suppl. methods). Protein separation and immunodetection were generated with the Jess™ automated capillary nano-immunoassay platform (Protein Simple, Bio-techne, 004-650) with chemiluminescence. Molecular mass marker (12-230 kDa), rabbit IgG anti-ApoE monoclonal (Abcam ab52607 [EP1374Y], 1 to 800 dilution) and anti-rabbit detection module (Novus DM-001) were loaded according to manufacturer instructions. Compass exposure setting 1 (1 second) was applied for visualization of peaks. Distinct peaks corresponding to molecular weights of ApoE monomers, dimers, trimers, tetramers and pentamers are designated as 1, 2, 3, 4 and 5, respectively. Incubation with either individual PUFA-PEs or a mixture of PUFA-PEs plus cholesterol increased ApoE3 dimerization and ApoE2 multimerization with the following apparent order of potency PE-P18:0/22:6n-3>PUFA-PE mixture plus cholesterol>PE-18:0/22:6n-3>PE-18:0/22:4n-6. Cholesterol and the synthetic phospholipid DMPC-14:0/14:0 had little or no effect on ApoE3 dimerization or ApoE2 multimerization. PUFA-phospholipid induced multimerization was mostly (but not completely) reversed by DTT. Capillary nano-immunoassay chromatograms are representative of experiments performed three times. Abbreviations: PUFA-PE, polyunsaturated fatty acid-containing ethanolamine phospholipid; DTT, dithiothreitol.

### 3.3. Which disulfide-linked ApoE-heteromeric complexes are present in human brain?

We posited that in human brain the presence of Cys endows ApoE3 and ApoE2 (but not ApoE4) with the ability to form heteromeric complexes with glia-secreted apolipoproteins containing free Cys residues such as ApoJ_2_ and ApoD (**Fig 3**).[39-41] An initial approach to search for these putative human-specific ApoE-ApoJ_2_ and ApoE-ApoD heteromers is to use anti-ApoE, anti-ApoD, and/or anti-ApoJ antibodies to immunoprecipitate lipoproteins from frozen human brain specimens under strict non-reducing conditions to prevent cleavage of intermolecular disulfide bridges. Pulled down material can be analyzed for the presence of disulfide-linked heteromers using western blot, with direct comparison of immunolabeled proteins under reducing and non-reducing conditions.

### 3.4. Do disulfide-linked ApoE homo- and heteromeric complexes protect lipid cargo from peroxidation?

Previous studies have demonstrated that ApoE3 possesses more antioxidant activity than ApoE4.[60, 61] Butterfield et al [62] hypothesized that aldehyde-scavenging activity of free Cys residues in ApoE3 and ApoE2 decrease AD risk. Notably, however, with only one or two Cys residues per 299 amino acids, ApoE3 and ApoE2 contain much lower proportions of Cys than established endogenous aldehyde-scavenging peptides such as glutathione.[63] Here we propose an alternative hypothesis wherein intact disulfide bridges (rather than free Cys) decrease AD risk specifically by inducing conformational changes that conceal and protect vulnerable PUFA-containing phospholipids from peroxidation (**Fig 2-4**). Guo et al. recently showed [64] that synthetic ApoE4- and cholesteryl arachidonate-containing lipoparticles increased neuronal lipofuscinosis and 4-hydoxynonenal (marker of lipid peroxidation), with deleterious effects of ApoE4 attributed to increased LDLR-mediated internalization of cholesteryl esters.[64] The ApoE-containing lipoparticles provided by Guo et al. did not contain PUFA-containing phospholipids, which are far more abundant than cholesteryl esters in human brain lipoproteins and parenchyma.[65, 66] Moreover, the ApoE-containing lipoparticles used by Guo et al. were treated with the thiol-reducing agents DTT and TCEP,[64] precluding assessment of the lipid-protecting role of intact intermolecular disulfide bridges posited here.

To test our hypothesis wherein the lipid-protecting effects of Cys in ApoE3 and ApoE2 are specifically due to the ability to form intermolecular disulfide bridges, one must first confirm whether there is a gradient of protective effects corresponding to abundance of Cys (ApoE2>ApoE3>ApoE4). To accomplish this, PUFA-phospholipid and ApoE-containing lipoparticles can be subjected to peroxidation and the extent of lipid peroxidation products can be quantified using established assays.[67, 68] Next, to determine whether protective effects are attributable to intact disulfide bridges, one can compare the extent of peroxidation in ApoE2 and ApoE3-containing dimers and multimers (with disulfide bridges intact) to their corresponding monomers lacking intact disulfide bridges. According to the present hypothesis, ApoE4 monomers (and ApoE2 and ApoE3 monomers without intact disulfide bridges) will be highly vulnerable to lipid peroxidation. By contrast, lipoparticles containing ApoE2 multimers and ApoE3 dimers with intact intermolecular disulfide bridges will be highly resistant and partially resistant to peroxidation, respectively.

### 3.5. Do intact disulfides in ApoE suppress Tau phosphorylation and Aβ secretion?

The next objective is to determine whether peroxidized ApoE4-containing lipoproteins induce neuronal GSK3-induced Tau phosphorylation, and/or Aβ secretion. To accomplish this, one must first generate and validate a human neuronal cell line that is capable of both Tau phosphorylation and Aβ secretion, then determine whether treatment with peroxidized ApoE4-containing lipoparticles evokes Tau phosphorylation and/or Aβ secretion compared to non-peroxidized ApoE4-containing lipoparticles. Quantitative assessment of pTau and Aβ can be accomplished using western blot with validated anti-pTau antibodies and commercially available ELISA-based assays, respectively. Having established that peroxidized ApoE4 lipoparticles induce Tau phosphorylation and Aβ secretion, the next objective would be to determine whether the disulfide bridges that are present in ApoE2 and ApoE3 multimers and dimers but absent in ApoE4 monomers (and ApoE2/ApoE3 monomers following cleavage), suppress Tau phosphorylation and Aβ secretion. This could be accomplished by treating neurons with peroxidized ApoE2, ApoE3, and ApoE4-containing lipoparticles. As noted above, when comparing ApoE2 and ApoE3 homo- and hetero-dimers and multimers, respectively, to their corresponding reduced monomers, great care should be taken to minimize or eliminate confounding effects of thiol-reducing agents.

## 4. Consequence of the hypothesis and discussion

### 4.1. Implications for mechanistic understanding of AD pathogenesis

The leading hypotheses proposed to explain the etiology of AD—the amyloid cascade and Tau prion-like propagation[69-72]—have limitations and inconsistencies, and are difficult to reconcile with one another and with other pathological observations and genetic risk factors for sporadic AD. In two recent publications,[15, 16] we provided extensive evidence supporting an alternative mechanistic paradigm and unifying model wherein the peroxidation of ApoE-containing lipoproteins and ensuing ApoE receptor-ligand disruption are the initiating molecular events that ultimately lead to AD in humans.[15, 16] Specifically, this work demonstrated that: (1) the same neurons that accumulate pTau in the earliest stages of AD strongly express ApoER2;[15, 16] (2) these same ApoER2-expressing neurons accumulate many other ApoER2-Dab1 pathway components;[15, 16] (3) lipid-peroxidation modified ApoE accumulates together with three native ApoER2 ligands (ApoE, Reelin, ApoJ) in extracellular Aβ-containing plaques that are surrounded by dystrophic neurites containing pTau and multiple other ApoER2-Dab1 pathway components (i.e. neuritic plaques);[15] and (4) Lys and His-enriched residues within the binding motifs of ApoE and its receptor ApoER2 are highly vulnerable to peroxidation, adduction, and crosslinking,[15] which were only partially reversible in an acidic milieu modeling the lysosome.[15] These collective findings, which strongly implicated peroxidation of ApoE-containing lipoproteins and ensuing ApoE receptor-ligand disruption in AD pathogenesis, have important implications for mechanistic understanding and therapeutic targeting of AD in humans.[15, 16] However, this mechanistic paradigm did not account for why *APOE* allele variants are the dominant genetic risk factors for sporadic AD (**Fig 1**). The present hypothesis—which attempts to fill that gap by proposing an unambiguous explanation for the isoform-dependent effects of *APOE* on lipoprotein peroxidation and AD risk—is integrated into our unifying model wherein peroxidation of ApoE-containing lipoproteins and consequent ApoE receptor-ligand disruption are the initiating molecular events that ultimately lead to AD in humans.[15, 16] This combined hypothesis is attractive because it provides a plausible, unambiguous molecular mechanism that:

1. can explain enduring genetic and neuropathological puzzles linking ApoE and lipid peroxidation to AD (including divergent effects of *APOE2* and *APOE4* alleles, accumulation of ApoE in the core of neuritic plaques, and evidence for lipid peroxidation in the earliest stages of AD);
2. is traceable directly back to the Cys⟶Arg exchanges that are the fundamental difference between major *APOE* variants;
3. can reconcile loss of function and gain of toxic function aspects of ApoE4;
4. can integrate AD-defining Aβ and pTau pathologies with other established AD pathologies including accumulation of ApoE, ApoJ, and endolysosomal dysfunction, and emerging AD pathologies such as the co-accumulation of Dab1, Reelin, pPSD95 and ApoER2 (reviewed in [15, 16]);
5. can help explain the strong genetic link between *CLU* variants and AD;
6. can help explain why aging and environmental triggers for lipid peroxidation predispose to sporadic AD (reviewed in [15]);
7. provides tangible leads for the prevention and treatment of AD in humans.

### 4.2. Evolutionary hypothesis: *APOE3* and *APOE2* help humans protect brain enriched PUFA-phospholipids from peroxidation

*Apoe4*, which encodes zero Cys residues, is the ancestral allele shared by all mammals except for humans.[73] *APOE4* homozygosity was the default human genotype until ≈200,000 years ago when emergence of the human-specific *APOE3* allele provided a single Cys addition at position 112 in the mature protein.[74] *APOE3* has since become the predominant allele in modern human populations. The *APOE2* allele emerged more recently,[73] adding a second Cys at position 158. Intriguingly, emergence and selection of *APOE3* appears to have coincided temporally with both the exploitation of coastal habitats that provide easy and reliable access to foods that are rich sources of DHA, and the rapid expansion of grey matter in the modern human brain.[75-77] DHA-containing phospholipids, which have a unique ability to regulate lipid raft dynamics and dendritic arborization,[78-80] are enriched within synaptic membranes. The brains of modern humans have more neurons, more complex dendritic arbors and a higher synapse-to-neuron ratio than the brains of mice and even monkeys.[81-83] Thus, marine foods may help support and maintain unique aspects of modern human brains (reviewed in [76]). However, because DHA-containing lipids are highly vulnerable to peroxidation,[18, 84] and post-mitotic neurons in modern brains must survive and function for 75 years or longer, humans may have a greater need to protect these vulnerable lipids than other mammals. We thus speculate that the human-specific presence of Cys, and ability and super-ability for ApoE3 and ApoE2 to form intermolecular disulfide bridges, provide advantages over ApoE4 in protecting brain-enriched PUFA-phospholipids from peroxidation.

### 4.3. Have rodent models obscured the crucial lipid-protecting role of disulfide-bridges in ApoE?

Rodent models that have been historically used for AD research are homozygous for ancestral *Apoe4* and therefore cannot form disulfide-linked homo- or heteromeric ApoE complexes. Newer, humanized APOE transgenic mouse models [85] enable ApoE homo-dimerization; however, the lack of unpaired Cys in rodent ApoD and ApoJ proteoforms precludes formation of disulfide-linked heteromeric ApoE-containing complexes such as those shown in **Fig 3**. Thus, reliance on rodent models for preclinical research—together with the use of thiol-reducing reagents and non-peroxidizable lipids such as DMPC for ApoE lipidation studies—may have obscured the crucial lipid-protecting role of disulfide bridges in ApoE-containing lipoproteins in the human brain.

### 4.4. Implications for AD therapeutics and prevention

Our mechanistic paradigm offers tangible predictions regarding the potential efficacy of existing AD therapeutics and clues for designing new therapeutics and preventive strategies. For example, our model predicts that *APOE2* gene therapy [86] will decrease AD risk in *APOE3* homozygotes by enhancing formation of lipid-protecting disulfide bridges between existing ApoE3 and newly-generated ApoE2 molecules. By contrast, since *APOE4* homozygotes lack Cys and therefore cannot form disulfide-linked ApoE2-ApoE4 complexes, our model predicts that such interventions will not affect gain of toxic function aspects of ApoE4, and thus provide only modest or no benefit in this high-risk population. This prediction is consistent with epidemiological evidence indicating that AD risk in *APOE2*/*4* heterozygotes is higher than *APOE3* homozygotes and nearly as high as *APOE3*/*4* heterozygotes (**Fig 1**). Therapeutics that lower ApoE4 [48] and therapeutics, dietary components, and preventive or environmental strategies that target underlying causes of lipoprotein peroxidation (reviewed in suppl appendix of [15]), are predicted to provide larger clinical benefits in *APOE4* carriers and homozygotes.

### 4.5. Summary and Conclusion

More than three decades have passed since the *APOE4* and *APOE2* alleles were identified as the strongest genetic driver and protective factor for AD, respectively; yet the specific molecular feature(s) and mechanisms underlying these isoform-dependent effects remain unknown. We propose a plausible, unambiguous solution to this enduring puzzle wherein the super-ability (for ApoE2), intermediate ability (for ApoE3) or inability (for ApoE4) to form lipid-protecting intermolecular disulfide bridges is the central molecular determinant accounting for the disparate effects of *APOE* variants on AD risk in humans. Veracity of this hypothesis has important implications for mechanistic understanding and therapeutic targeting of AD in humans, and as such, warrants rigorous investigation.

## List of abbreviations

1-1(Z)-Octadecenyl-2-Docosahexaenoyl-sn-glycero-3-PE (PE-P18:0/22:6n-3)

1-Stearoyl-2-Docosahexaenoyl-sn-glycero-3-PE (PE-18:0/22:6n-3)

1-Stearoyl-2-Docosatetraenoyl-sn-glycero-3-PE (PE-18:0/22:4n-6)

Aβ: amyloid-β
AD: late-onset, sporadic Alzheimer’s disease
ApoE: Apolipoprotein E
ApoER2: ApoE receptor 2
Arg: arginine (also designated R)
Cys: cysteine (also designated C)
Dab1: Disabled homolog-1
DMPC: 1,2-Dimyristoyl-sn-glycero-2-PC (PC-14:0/14:0)
DHA: docosahexaenoic acid
DTT: dithiothreitol
HNE: hydroxynonenal
LDLR: low density lipoprotein receptor
Lys: lysine (also designated K)
MCI: mild cognitive impairment
Met: methionine
PE: phosphatidylethanolamine
PC: phosphatidylcholine
PSD95: postsynaptic density-95
pPSD95: phosphorylated PSD95
pTau: hyper-phosphorylated Tau
PUFA: polyunsaturated fatty acid
Tau: microtubule associated protein tau
TCEP: tris(2-carboxyethyl) phosphine

## Acknowledgements

We are grateful to Marc Raley for graphic art in Figs 1-3, Mark Horowitz for endnote referencing and assistance with manuscript submission, Josephine Egan and Qing Liu for proofreading, and Daisy Zamora for statistical interpretation in genetics research.

## Author contributions

Conceptualization, CER; literature review, CER, RGC; experiments, GSK, XL, CER; resources, CER; writing—original draft, CER; writing—review and editing, GSK, RGC, XL. All authors read and agreed to the published version of the manuscript.

## Funding

This project was supported by the intramural programs of the National Institute on Aging (1ZIAAG000453) and the National institute on Alcohol Abuse and Alcoholism, National Institutes of Health.

## Availability of data and materials

Data available from the corresponding author on reasonable request.

## Declarations

No competing interests to report.

## Notes

### Competing Interest Statement

The authors have declared no competing interest.

