## Supplement for "Lipid-protecting disulfide bridges are the missing molecular link between ApoE4 and sporadic Alzheimer’s disease in humans"

### Supplementary methods

#### Testing effects of lipids on disulfide-dependent ApoE dimer/multimerization

The recombinant human ApoE proteins and lipids that were used for incubations are listed the in the **Table S1**. To generate synthetic ApoE-containing lipoparticles, recombinant ApoE has traditionally been lipidated with peroxidation-resistant lipids such as 1,2-Dimyristoyl-sn-glycero-3-phosphatidylcholine (DMPC-14:0/14:0). Although DMPC is an artificial lipid, it was presumably selected to limit the extent of lipid degradation, peroxidation, and subsequent ApoE aggregation during a multi-step lipidation and purification procedures. Lipidation of ApoE has also typically included the addition of sodium cholate (a bile acid produced in liver) and thiol-reducing agents such as dithiothreitol (DTT), tris(2-carboxyethyl) phosphine (TCEP), and/or β-mercaptoethanol. Since DMPC is resistant to peroxidation and thiol-reducing agents cleave disulfide bonds, use of these reagents precluded evaluation of our hypothesis wherein intact disulfide bridges perform a crucial function by concealing and protecting vulnerable PUFA-containing phospholipids from peroxidation.

To investigate whether exposure to PUFA-containing phospholipids induces disulfide bridge formation in ApoE, we developed a simplified method for incubating lipids with ApoE that was designed to limit PUFA degradation, ensuing peroxidation, and ApoE aggregation, without addition of thiol-reducing agents, DMPC, or any other extraneous reagents. Briefly, 10 ug of recombinant lyophilized ApoE protein (ProSci: ApoE2 [40-140], ApoE3 [40-136], ApoE4 [40-138]) was diluted in PBS and vortexed for 30s. Equimolar amounts of brain-enriched 1-1(Z)-Octadecenyl-2-Docosahexaenoyl-sn-glycero-3-PE (PE-P18:0/22:6n-3), 1-Stearoyl-2-Docosahexaenoyl-sn-glycero-3-PE (PE-18:0/22:6n-3), 1-Stearoyl-2-Docosatetraenoyl-sn-glycero-3-PE (PE-18:0/22:4n-6), cholesterol, or DMPC in ethanol were added to ApoE and the mixture was incubated for 2 h at 37C, with vortexing for 30s every 30 minutes. Amounts of lipids used for the experiments reported in the main paper were selected based on the results of pilot dose-response studies, which revealed that an 8 to 1 molar ratio of PE-P18:0/22:6n-3 to ApoE was sufficient to markedly increase ApoE2 dimer/multimerization (**Fig S1**). By contrast, the peroxidation-resistant lipid DMPC and cholesterol had no appreciable effect even up to 43 to 1 molar ratio.

#### Protein separation and immunodetection

ApoE protein separation and immunodetection were generated with the Jess™ automated capillary nano-immunoassay platform (Protein Simple, Bio-techne, 004-650) using 25 column Jess™ plates with chemiluminescence. Molecular weight marker (12-230 kDa), the rabbit anti-ApoE monoclonal (Abcam ab52607 [EP1374Y], 1 to 800 dilution) and the anti-rabbit secondary HRP-conjugated antibody (Novus DM-001) were loaded according to manufacturer instructions. Compass exposure setting 1 (1 second) or high dynamic range settings were applied to optimize visualization of peaks corresponding to molecular weights of ApoE monomers, dimers, trimers, tetramers and pentamers. To assess the impact of disulfide bridges, protein separation and immunodetection assays were completed both with and without the addition of DTT.

Supplementary materials for Lipid-protecting disulfide bridges are the missing molecular link between ApoE4 and sporadic Alzheimer's disease in humans (Ramsden et al. 2025)

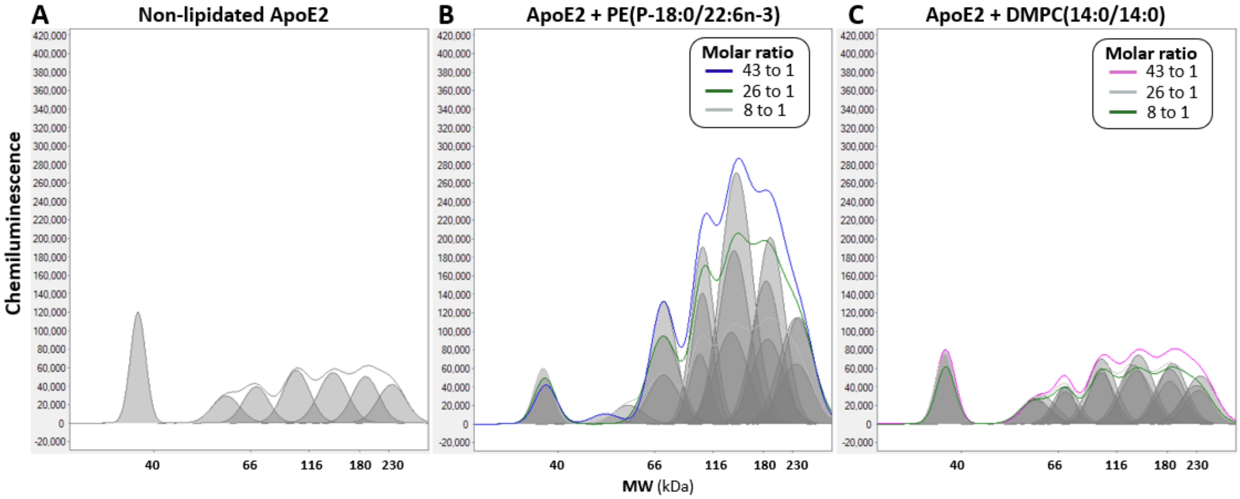


***Figure S1. Dose-dependent effects of PE-(P18:0/22:6n-3) and DMPC-(14:0/14:0) on disulfide-dependent ApoE2 multimerization***

*Recombinant ApoE2 protein was incubated with: (**A**) no added lipid, (**B**) PE-(P18:0/22:6n-3), or (**C**) DMPC-(14:0/14:0), with lipid to ApoE molar ratios of 8:1, 26:1, and 43:1. The Jess$^{TM}$ automated capillary nano-immunoassay platform (Protein Simple, Bio-techne, 004-650) with chemiluminescence was used for protein separation and immunodetection. Molecular weight marker (12-230 kDa), rabbit IgG anti-ApoE monoclonal (Abcam ab52607 [EP1374Y], 1 to 800 dilution) and anti-rabbit detection module (Novus DM-001) were loaded according to manufacturer instructions with the Compass high dynamic range exposure setting applied for visualization of peaks. Incubation with PE-(P18:0/22:6n-3) markedly increased ApoE2 multimerization in a dose-dependent manner (**B**), with substantial effects evident even with an 8:1 molar ratio. By contrast, the artificial, peroxidation-resistant lipid DMPC had little or no effect on ApoE2 multimerization (**C**). Capillary nano-immunoassay chromatograms are representative of experiments performed three times.*

### Table S1. Key Resources

| *Reagent type or resource* | *Molecule or target* | *Designation* | *Type or source* | *Source* | *Catalog #* |
|---|---|---|---|---|---|
| **Antibodies** | | | | | |
| Antibody (primary) | Apolipoprotein E | ApoE | Rabbit IgG [EP1374Y] | Abcam | ab52607 |
| **Recombinant ApoE proteins** | | | | | |
| protein | Human recombinant ApoE2 | ApoE2 | E. Coli | ProSci | 40-140 |
| protein | Human recombinant ApoE3 | ApoE3 | E. Coli | ProSci | 40-136 |
| protein | Human recombinant ApoE4 | ApoE4 | E. Coli | ProSci | 40-138 |
| **Lipids** | | | | | |
| Lipid (artificial) | 1,2-Dimyristoyl-*sn*-glycero-3-PC | DMPC-(14:0/14:0) | Artificial, peroxidation-resistant phosphatidylcholine | Cayman Chemical | 15097 |
| Lipid (brain-enriched) | Non-esterified Cholesterol | Chol | Free form of cholesterol | Cayman Chemical | 39088 |
| Lipid (brain-enriched) | 1-1(Z)-Octadecenyl-2-Docosahexaenoyl-sn-glycero-3-PE | PE-(P18:0/22:6n-3) | PUFA-PE | Cayman Chemical | 37138 |
| Lipid (brain-enriched) | 1-Stearoyl-2-Docosahexaenoyl-sn-glycero-3-PE | PE-(18:0/22:6n-3) | PUFA-PE | Cayman Chemical | 41382 |
| Lipid (brain-enriched) | 1-Stearoyl-2-Adrenoyl-*sn*-glycero-3-PE | PE-(18:0/22:4n-6) | PUFA-PE | Cayman Chemical | 33661 |

Abbreviations: PUFA, polyunsaturated fatty acid; PC, phosphatidylcholine; PE, phosphatidylethanolamine
